# Ubiquitination of the yeast receptor Atg32 modulates mitophagy

**DOI:** 10.1101/652933

**Authors:** Pierre Vigié, Cécile Gonzalez, Stephen Manon, Ingrid Bhatia-Kissova, Nadine Camougrand

## Abstract

Mitophagy, the process that degrades mitochondria selectively through autophagy, is involved in the quality control of these organelles. In yeast, the presence of the Atg32 protein on the outer mitochondrial membrane allows for the recognition and targeting of superfluous or damaged mitochondria for degradation. Some posttranslational modifications, such as phosphorylation, are crucial for the execution of the mitophagy process. In our study, we showed that in the stationary phase of growth, and to a lesser extent during starvation, the Atg32 protein level decreases. The fact that a decline in Atg32 level can be prevented by inhibition of the proteolytic activity of proteasome may indicate that Atg32 is also ubiquitylated. In fact, mass spectrometry analysis of purified Atg32 protein showed ubiquitination of lysine residue in position 282. These different patterns of posttranslational modifications of Atg32 could allow cells to control the mitophagy process carefully.

## Introduction

Mitochondria are organelles in charge of many crucial functions in cells. In eukaryotes, they are known to be the powerhouse of cells by producing adenosine triphosphate via oxidative phosphorylations; they are also involved in different synthesis pathways such as synthesis of certain amino acids and lipids and in signaling pathways such as apoptosis and calcium. Mitochondrial dysfunctions have been associated with aging and with an increasing number of pathologies including neurodegenerative diseases, cancer, and metabolic disturbances. To maintain cell survival and homeostasis, cells developed ways to remove superfluous or damaged cell components and organelles such as mitochondria. Mitophagy is one the processes involved in the mitochondrial quality control. Mitophagy is a selective form of autophagy.^1,2^ Macroautophagy (hereafter called autophagy) is conserved among eukaryotic species and involves specific lytic compartments: the vacuole in yeast and lysosomes in mammals. Autophagy involves specific autophagy-related (Atg) proteins. To date, more than 40 Atg proteins have been characterized; about half of them are involved in the core autophagy machinery and the rest is implicated in specific autophagy processes and regulation. In yeast, autophagy is initiated in the pre-autophagosomal structure (PAS) located next to the vacuole. The PAS comprises proteins and lipids constantly in motion. The phagophore, a double phospholipid membrane structure, emerges from the PAS and sequesters cytosolic components in vesicles called autophagosomes. Autophagosomes merge with the vacuolar membrane, releasing the trapped cell components into the lumen, where hydrolases degrade them. In yeast, Atg32 is the only characterized mitophagy receptor, which suggest mitophagy could be a simpler mechanism in yeast than in mammals. Atg32 protein was characterized by 2 teams in 2009 that aimed to find proteins involved in mitophagy specifically.^3,4^ Atg32 is an outer mitochondrial membrane protein with its C-terminus in the intermembrane space and its N-terminus in the cytosol. Atg32 expression increases in cells during respiratory growth. When mitophagy is induced, Atg32 interacts with both Atg8, a protein anchored to phagophore and autophagosome membranes due to a PE moiety, and Atg11, a scaffold protein required for other selective autophagy processes.^3,4,5,6^ Both interactions are required for the recruitment of mitochondria to the phagophore followed by their sequestration within autophagosomes and their degradation in the vacuole.

To date, several post-translational modifications of Atg32 were described. Protein is phosphorylated on serines 114 and 119 by Ck2 kinase during mitophagy induction, and serine 114 phosphorylation mediates Atg32-Atg11 interaction.^5,7,8^ Conversely, under growing conditions, Ppg1 is responsible for dephosphorylation of Atg32 and mitophagy inhibition.^9^ Mao et al. (2011) also showed that other kinases involved in mitogen-activated protein kinase pathways such as Slt2, Hog2, and Bck1 regulate mitophagy^10^. Yme1 is responsible for a partial Atg32 C-terminus processing, in the intermembrane space when mitophagy is triggered.^11^ Atg32 protein expression is also regulated by intracellular reduced glutathione.^12^ Indeed, the absence of an Opi3 protein, an enzyme required in conversion of phosphatidylethanolamine to phosphatidylcholine, causes an increase in intracellular reduced glutathione leading, to Atg32 protein expression decrease.^12^ Moreover, the absence of some components of NatA N-acetyltransferase complex also results in a decrease in Atg32 protein expression. However, NatA’s precise role concerning Atg32 protein is still unclear.^13^ More recently, Levchenko et al. (2016) detected another post-translational modification of Atg32 when mitophagy was induced by rapamycin treatment in a *Δpep4* background where vacuolar proteolysis is impaired.

However, its precise nature remains to be characterized.^14^ Xia et al. (2018) have examined the structure of Atg32 and have identified a structured domain in Atg32 that is essential for the initiation of mitophagy because it is required for the proteolysis of the C-terminal domain of Atg32 and the subsequent recruitment of Atg11.^15^ The solution structure of this domain was determined by NMR spectroscopy, revealing that Atg32 contains a previously undescribed pseudo-receiver (PsR) domain, within the cytosolic region of Atg32 comprising residues 200-341. Their data suggest that the PsR domain of Atg32 regulates Atg32 activation and the initiation of mitophagy.^15^ All these data showed that the regulation of Atg32 protein is complex.

In this work, we explored the existence of an interplay between mitophagy and proteasome in yeast. We demonstrated a novel post-translation modification of Atg32, ubiquitination at least at Lysine 282 residue, which is involved in regulation of Atg32 turnover and its function in mitophagy.

## Results

### Expression of the mitophagy receptor Atg32 decreases during stationary phase

Wang et al. (2013) found that when mitophagy is induced by nitrogen starvation, Atg32 C-terminus, located in the IMS, is processed by Yme1, an i-AAA protease found in the inner mitochondrial membrane.^11^ Atg32 might be synthesized as an immature form of the protein and processed on the mitochondrial surface so that specific mitochondria could be degraded selectively. Moreover, when cells were grown on a respiratory carbon source like lactate before undergoing to nitrogen starvation, this cleavage is insignifiant (i.e. 5%) compared to what happens in glucose grown cells. This cleavage is also dependent on the promoter used to express the Atg32 protein.^11^ In their paper, Wang et al. did not investigate the behaviour of the Atg32 protein during the stationary phase of growth. To study Atg32 expression in cells grown in a lactate-containing medium, we C-terminally tagged the Atg32 protein, under its own promoter, with a V5 epitope tag (Atg32-V5). Atg32 expression was determined from cells harvested in the mid-exponential phase of growth (T0), 8 hours later (8h), and after one day (24h) and 2 days (48h) of culture. We found that under respiratory conditions, the Atg32 protein level is decreased during cell growth and almost completely disappeared in the stationary phase of growth (Fig. 1A,B). We observed that when the Atg32 protein is N-terminally tagged with an HA epitope, its expression also decreased during the stationary growth phase (Fig. S1A,B). At the same time, during the first hours of nitrogen starvation, Atg32-V5 levels did not vary much (Fig. 1C,D). We ensured the fusion protein Atg32-V5 was localized to mitochondria (Fig. 2A) and was able to reverse the mitophagy defect of the *atg32Δ* mutant (Fig. 2B,C). These observations are in accordance with the data of Levchenko et al., who showed that C-terminal tagging of Atg32 by a ZZ tag did not interfere with mitophagy induced by nitrogen starvation or rapamycin treatment.^14^ Moreover, we observed in *atg5Δ, atg8Δ*, and *atg11Δ* mutants the same level of Atg32 at T0 and a decrease in the stationary phase of growth (Fig. S2A,B). Okamoto et al saw a similar disappearance of Atg32 in *atg7Δ* mutant.^4^ Assuming that Atg32 is only essential for mitophagy, this finding was surprising. The reason for this finding is not certain; however, it is possible that a small quantity of Atg32 might be sufficient to mediate selective elimination of mitochondria. Alternatively, yet unidentified posttranslational modification of Atg32 might function in mitophagy. Also, we can not exclude the possibility that, during respiratory growth, Atg32 might be involved in yet unknown pathway(s) related to mitochondrial function(s).

**Figure 1:**
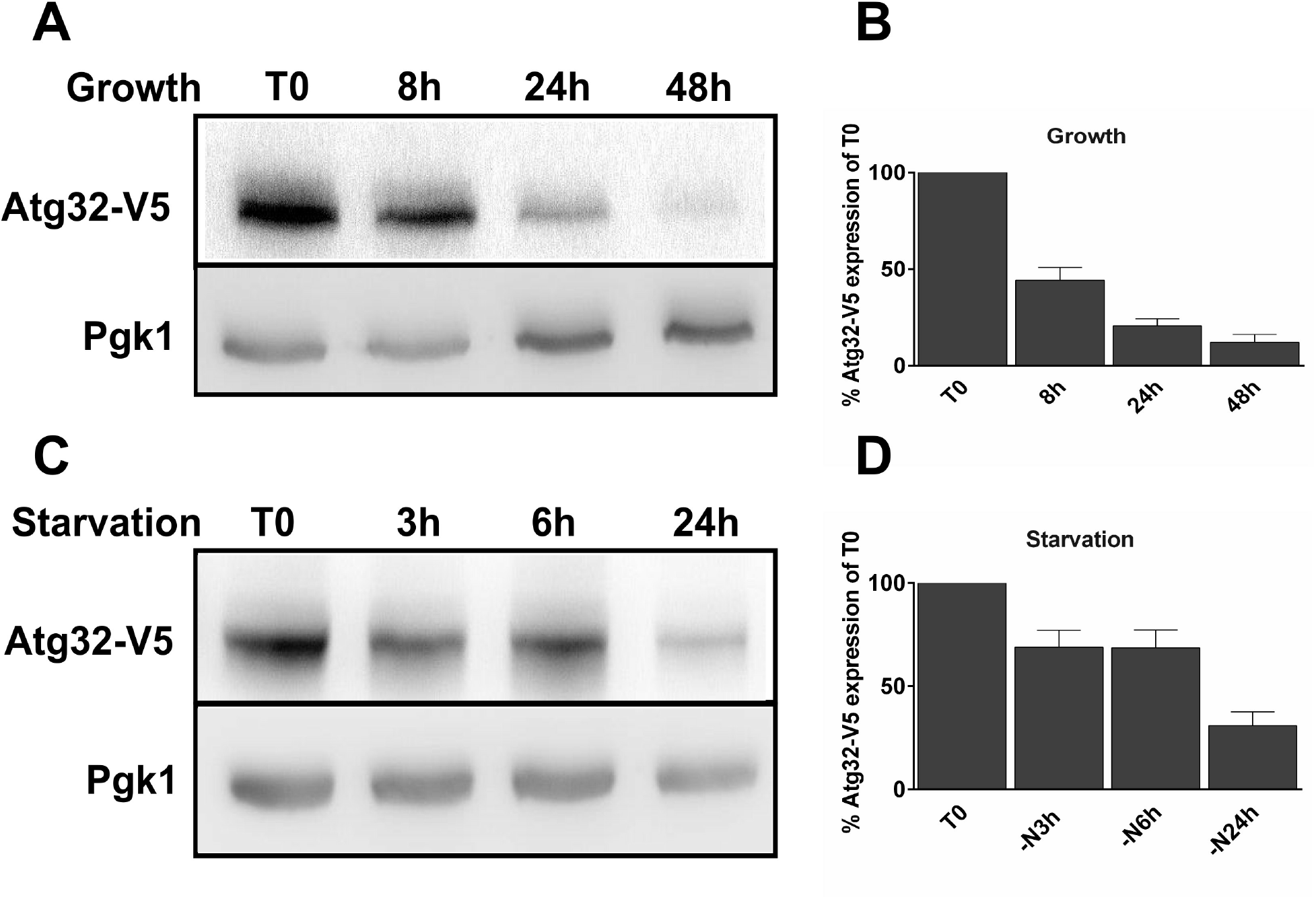
Expression of Atg32-V5 protein decreases during cell growth. *atg32Δ* mutant cells grown in a lactate-containing medium and expressing Atg32-V5 recombinant protein were harvested in a mid-exponential phase of growth (T0) and after 8-, 24- and 48 hours from T0 (**A**) or after 3-, 6- or 24 hours of nitrogen starvation (**C**). Total protein extracts were prepared afterwards and protein samples were analyzed by western blots. Anti-V5 antibody was used to visualize Atg32-V5 recombinant protein, and Pgk1, the cytosolic phosphoglycerate kinase, was used as loading control. (**B,D**) Atg32-V5 expression was quantified as the percentage of Atg32-V5 level of T0 (100%). Three independent experiments were carried out for each conditions.

**Figure 2:**
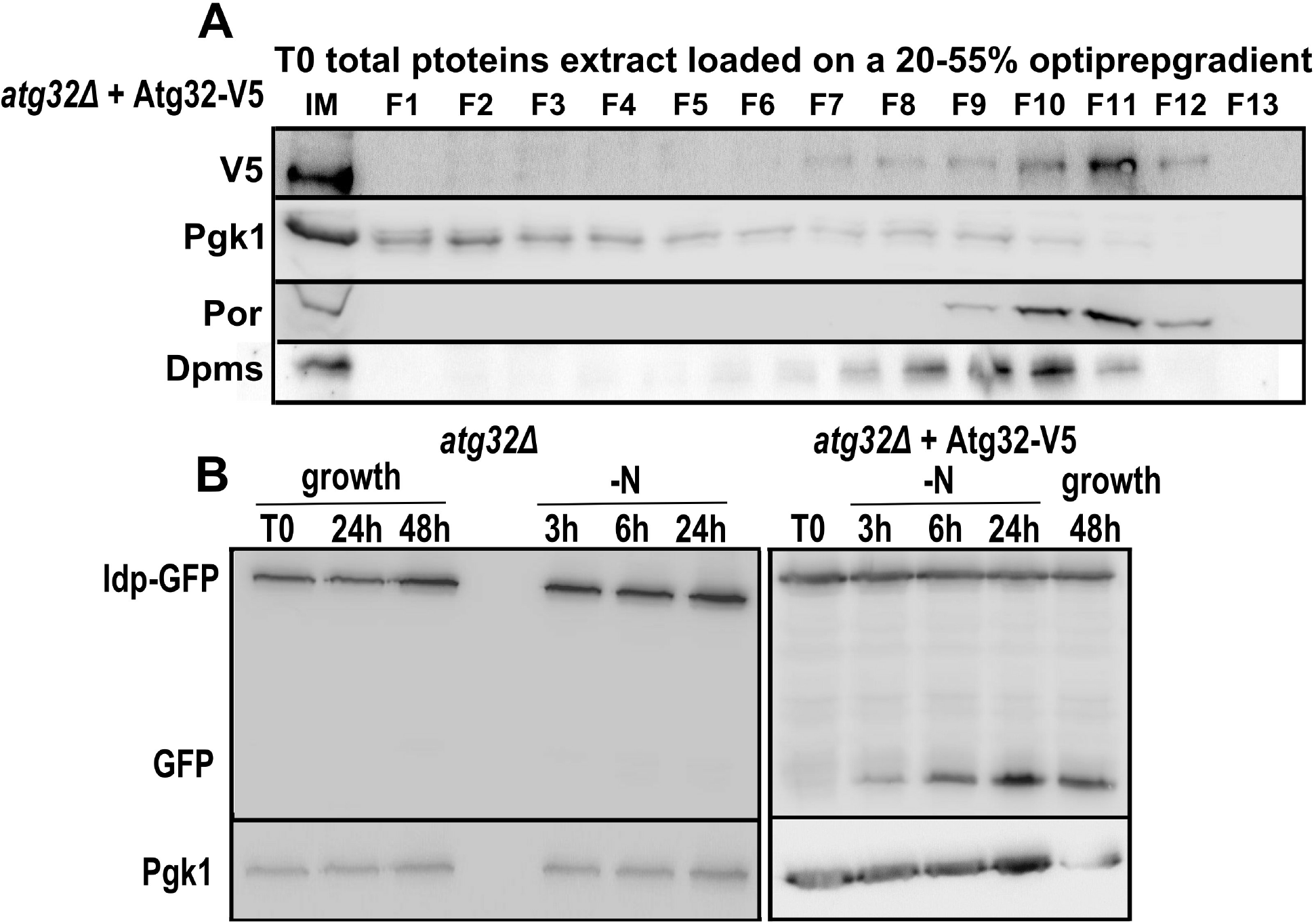
Atg32-V5 protein localizes to mitochondria and restores mitophagy in *atg32Δ* mutant strain. **(A)** *atg32Δ* mutant cells grown in a medium supplemented with a respiratory carbon source lactate and expressing Atg32-V5 fusion protein were harvested in a midexponential phase of growth. Then, cells were lysed and cell lysates were separated in a 20-55% OptiprepTM density gradient. Fractions were collected after centrifugation and analyzed by western blots. Anti-V5 antibody was used to visualize Atg32-V5 and anti-Por1 to detect mitochondria-containing fractions. Pgk1 was used as a cytosolic marker and Dpms as a ER marker. Each experiment was performed twice. **(B)** Mitophagy was assessed using Idp1-GFP tool in *atg32Δ* cells or *atg32Δ* expressing Atg32-V5. Cells were harvested in an exponential phase of growth (T0), or after 3 hours (-N3h), 6 hours (-N6h) and 24 hours (-N24h) of nitrogen starvation or during a course of cell growth (24 or 48 hours). The corresponding total protein extracts were separated by SDS PAGE and analyzed by western blots using anti-GFP antibody. Pgk1, the cytosolic phosphoglycerate kinase, was used as loading control. Results were obtained from 3 independent experiments.

### Effect of proteasome inhibition on Atg32 expression

What affects the expression of Atg32? To understand why the Atg32 level decreases when mitophagy is induced, we evaluated the effect of inhibitors of the two main cellular proteolytic pathways (ubiquitin-proteasome and vacuolar-autophagy system) on the Atg32 level. First, we observed that the addition of MG-132, a proteasome inhibitor, which reduces the degradation of ubiquitin-conjugated proteins in mammalian cells and effectively blocks the proteolytic activity of the 26S proteasome in yeast, counteracts Atg32 protein loss observed during the stationary growth phase (Fig. 3A,B), the first hours of nitrogen starvation (Fig. 3C,D), and rapamycin treatment (Fig. S3A). In figure 3A,C (lower panels), accumulation of ubiquitin-conjugated complexes (Ub) becomes easily detectable as they are not degraded in the presence of this proteasome inhibitor. Also, the Atg32-V5 level was restored with MG-132 treatment in *atg5Δ* mutant in the stationary phase of growth (Fig. S2C). The same behaviour was observed in the BY4742 strain expressing Atg32-V5 (Fig. S3B,C).To answer the question of whether the reduction of Atg32 protein level during the stationary phase in respiratory conditions is due to a decrease in Atg32 synthesis or an increase in Atg32 degradation, we examined *ATG32* promoter activity. In our study, we used a construct pPROM-*ATG32-β-galactosidase*, in which a reporter gene lacZ was expressed from the *ATG32* promoter. Interestingly, we found that *ATG32* promoter activity increased through the course of cell growth (Fig. 3E). In the stationary phase of growth, Atg32 expression was increased 3-to 4-fold compared with cells in an early exponential phase of growth (T0). MG-132 treatment prevented this rise in *ATG32* promoter activity. Together, our data showed, that under respiratory conditions, the Atg32 protein level decreased during mitophagy while *ATG32* promoter activity increased (and conversely), suggesting that the diminition of the Atg32 protein level during the stationary phase is due to an increase of Atg32 degradation. It is tempting to speculate that Atg32 might regulate its own level and activity.

**Figure 3:**
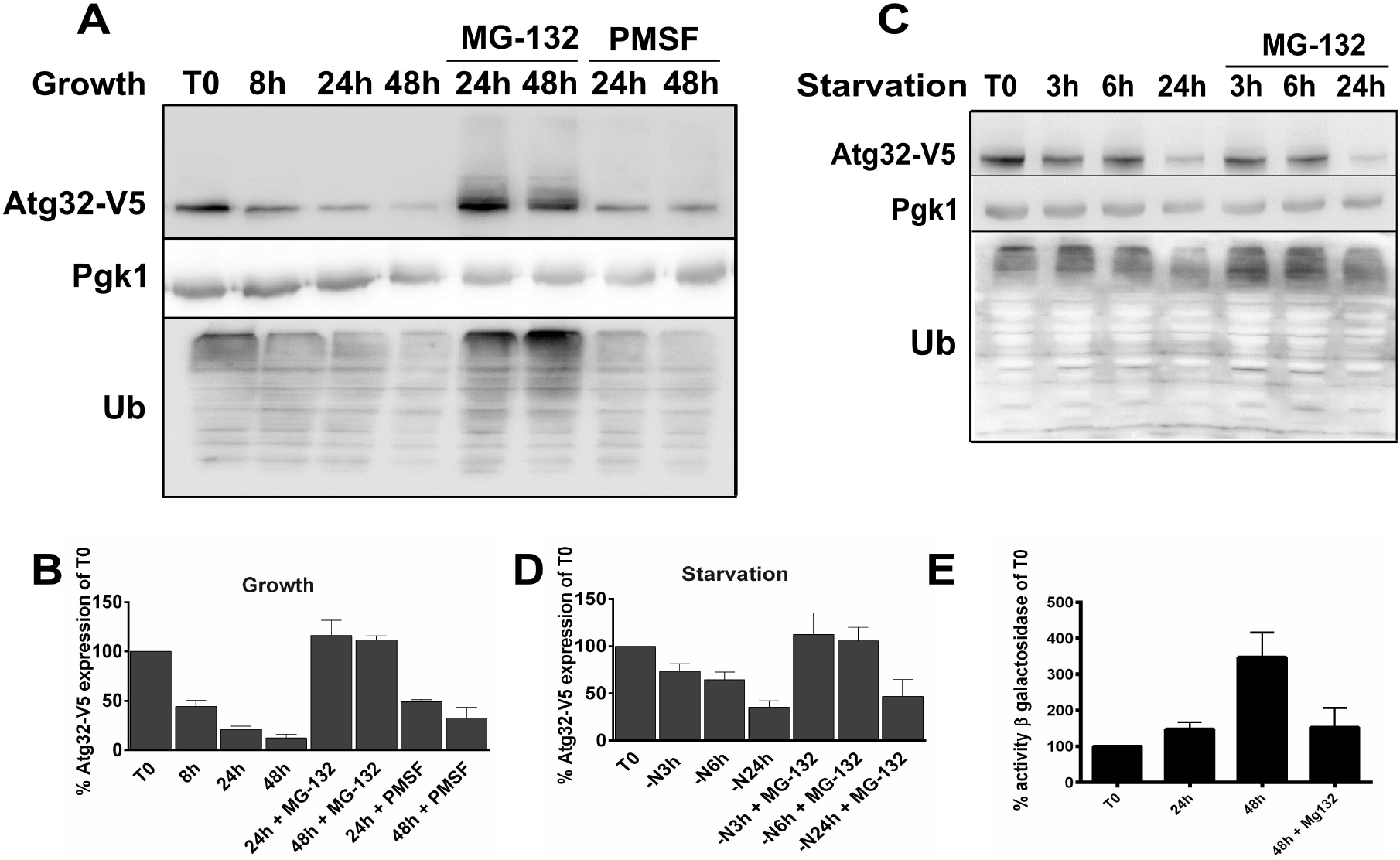
Decrease in the Atg32 protein level is prevented by inhibition of the proteolytic activity of proteasome. *atg32Δ* cells grown in a lactate-containing medium and expressing Atg32-V5 recombinant protein were harvested at different time points: **(A)** T0 that represents a mid-exponential phase of growth and then after 8-, 24- and 48 hours of cell growth or **(C)** after 3-, 6- or 24 hours of nitrogen starvation. To inhibit proteasome, 75 μM MG-132 + 0,003 % SDS were added to the cell culture at 8h time point. To inhibit vacuolar proteolysis, 2mM PMSF was added to the cell culture at 8h time point. Total protein extracts were prepared afterwards and protein samples were analyzed by western blots. Anti-V5 antibody was used to visualize Atg32-V5 recombinant protein; Pgk1, the cytosolic phosphoglycerate kinase, was used as loading control. Anti-ubiquitin (Ub) was used to detect the level of ubiquitinated proteins. **(B,D)** Atg32-V5 expression was quantified as the percentage of Atg32-V5 level of T0 (100%). Three independent experiments were carried out for each condition. **(E)** BY4742 cells grown in a lactate-containing medium and expressing β-galactosidase under the *ATG32* promoter were harvested in a mid-exponential phase of growth (T0) and after 24- and 48 hours from T0 in absence or presence of MG-132. After cells lysis, the β-galactosidase activity was measured. Results were obtained from 6 independent experiments and are expressed as the % of T0.

Meanwhile, addition of PMSF, an inhibitor of serine proteases, that blocks a number of yeast vacuolar proteases, such as proteinase B and carboxypeptidase Y, but does not affect proteasome function, only slightly rescued Atg32 protein levels, probably corresponding to the part of protein that is degraded by mitophagy (Fig. 3A). Consistent with these observations, it is possible that Atg32 is degraded in both autophagy-dependent and autophagy-independent manner. It should be noted that neither the addition of MG-132 nor PMSF significantly affected cell growth or growth yield (Fig. S4A). Furthemore, simultaneous addition of MG-132 and PMSF had no significant cumulative effect on reduction of the Atg32 levels (Fig. S4B).

The fact that treatment with MG-132 prevented the disappearance of Atg32 remarkably can indicate that Atg32 protein degradation seems to be directly or indirectly dependent on the proteasome activity. These results indicate that Atg32 is ubiquitinated and targeted for proteasomal degradation. Indeed, several bands with the molecular weight shift of Atg32-V5 in Figure 3A after MG-132 treatment (upper panel) during cell growth in a respiratory conditions may correspond to ubiquitinated Atg32 protein forms. At the same time, it does not exclude the possibility that some longer bands can (also) be phosphorylated or otherwise modified.

We next wanted to test what happens to the Atg32 levels in a proteasome mutant. Hence, we investigated the *pre2-2* mutant strain that has a severe defect in proteolysis mediated by the 26S proteasome. It was not able to grow in the lactate-containing medium, which we used in our study. For that reason, we grew the *pre2-2* mutant in a galactose-containing medium. It should be noted that mitophagy in a galactose-containing media is induced to a lesser extent compared to that one in lactate (Camougrand, not published results). Nevertheless, likewise in a lactate-containing medium, expression of the fusion protein Atg32-V5 decreased ostentatiously in the stationary phase of growth in medium with galactose in the control strain, whereas less pronounced change was observed for the *pre2-2* mutant, indicating the involvement of proteasome in the disappearance of Atg32-V5 protein during the stationary growth phase (Fig. 4 A,B).

**Figure 4:**
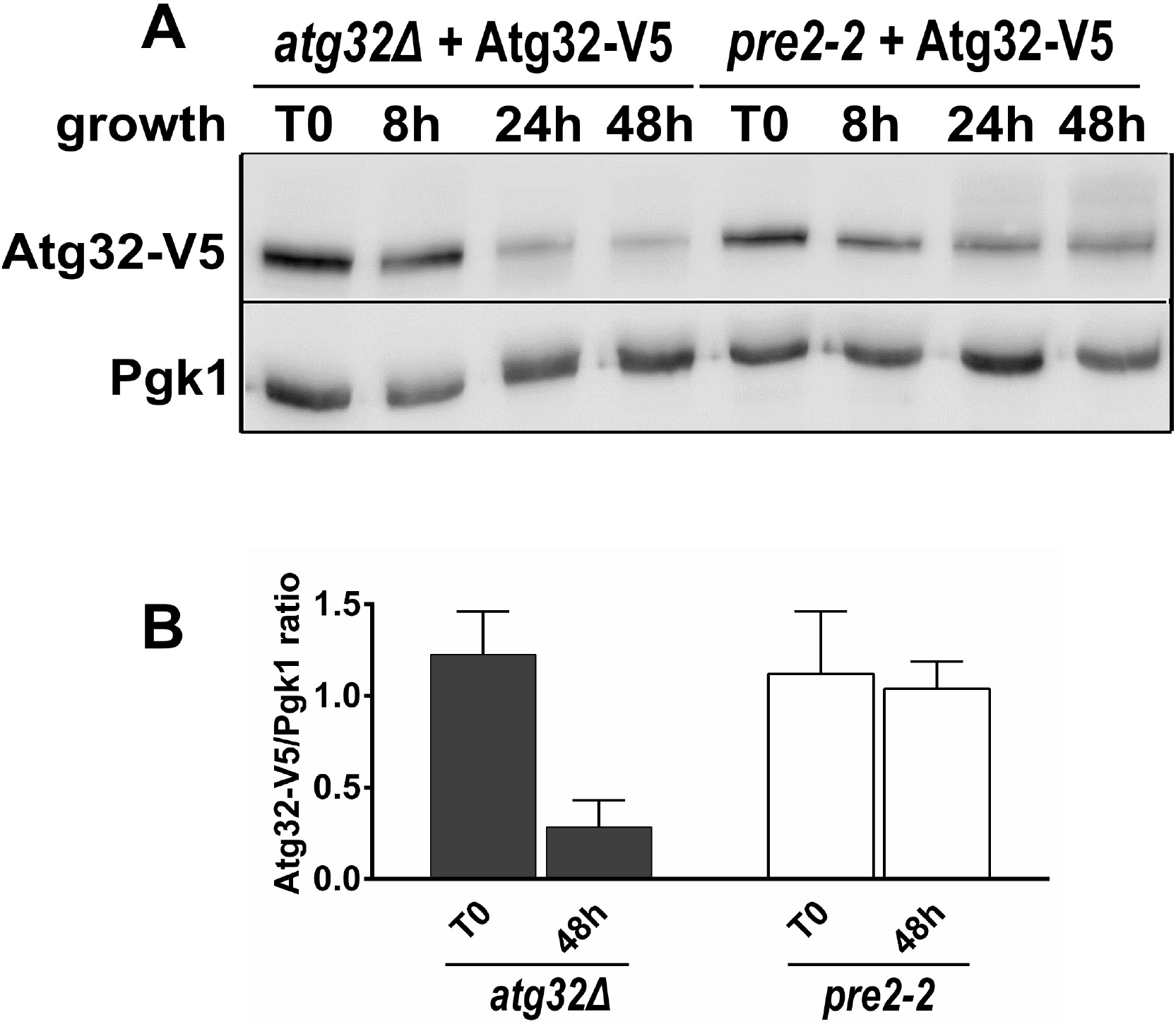
Study of Atg32-V5 expression in pre2-2 mutant. **(A)** atg32*Δ* and *pre2-2* mutant cells transformed with a plasmid expressing Atg32-V5 were grown in a minimal synthetic medium supplemented with 2% galactose as a carbon source. Cells were harvested in an early exponential phase of growth (T0) and after 8-, 24-, and 48 hours of cell growth. Total protein extracts were prepared afterwards and protein samples were analyzed by western blots. Anti-V5 antibody was used to visualize Atg32-V5 recombinant protein and Pgk1, the cytosolic phosphoglycerate kinase, was used as loading control. **(B)** Atg32-V5/Pgk1 ratios were quantified at T0 and T48 for all tested strains. Three independent experiments were carried out for each condition.

### Effect of proteasome inhibition on mitophagy

Previous research has shown that cells lacking Atg32 protein are deficient in mitophagy induction and that Atg32 overexpression is responsible for an increase in mitophagy.^3,4^ These published data suggest that mitophagy levels are correlated with the amount of Atg32. We observed that the Atg32-V5 protein did not disappear in the stationary phase of growth when cells were treated with MG-132. To check whether the higher protein level in the stationary phase of growth obtained after MG-132 treatment was correlated with an increase of mitophagy, we assessed mitophagy induction in this stage among cells treated or not treated with MG-132. We first studied mitophagy by western blots in cells expressing Idp1-GFP fusion protein. Cells were grown in respiratory conditions and harvested at different time points during growth in the presence or absence of MG-132. In cells without any treatment, the GFP band appeared in the early stationary phase (24h), suggesting that mitophagy was induced (Fig. 5A). At the same time points, in cells treated with MG-132, GFP bands were more intense, suggesting that proteasome inhibition results in an increase in mitophagy in the stationary phase of growth. To confirm the results we obtained by western blots, alkaline phosphatase (ALP) activity was measured in cells expressing mitochondria-targeted Pho8Δ60 (mtPho8Δ60) protein. In control cells, we observed a 2-fold increase in ALP activity in the stationary phase of growth (48h) compared to the midexponential phase of growth (T0; Fig. 5B). However, at the 48h time point with MG-132 treatment, we measured a 4-fold increase in ALP activity compared to T0, confirming that proteasome inhibition by MG-132 causes an increase in mitophagy in the stationary phase of growth compared to untreated cells. Notably, MG-132 had no stimulating effect on macroautophagy (data not shown).

**Figure 5:**
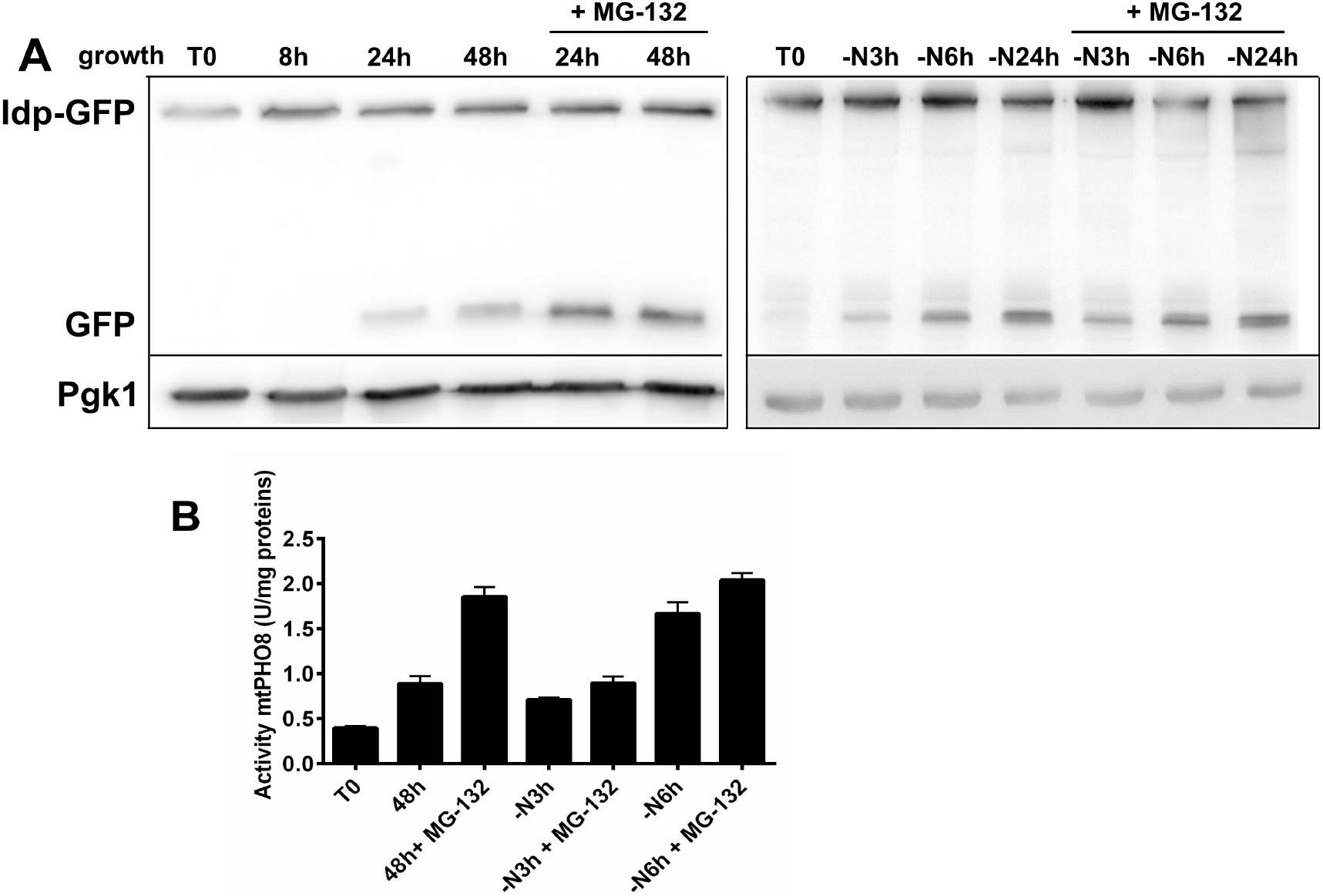
Inhibition of the proteolytic activity of proteasome increases mitophagy. **(A)** *atg32Δ* mutant cells expressing Atg32-V5 and Idp1-GFP proteins grown in a minimal synthetic medium with lactate were harvested at different times of growth: at T0 corresponds to a midexponential phase of growth and then after 8-, 24- and 48 hours of cell growth, or after 3-, 6- and 24 hours of nitrogen starvation. MG-132 (75 μM) and 0,003% SDS were added at the end of an exponential phase (time point 8h). In starved cells, MG-132 was added at the beginning of starvation. Total protein extracts from 2 × 10^7^ cells were prepared and separated on a 12,5% SDS-PAGE gel as described in the Material and methods section. Proteins were detected using antibodies against GFP (Roche) or Pgk1 (Invitrogen). **(B)** *atg32Δ* mutant cells expressing Atg32-V5 and mtPHO8Δ60 proteins grown in a minimal synthetic medium with lactate were harvested at different times of growth: at T0 that corresponds to a mid-exponential phase of growth and after 48 hours in the stationary phase of growth or after 3 and 6 hours of nitrogen starvation. MG-132 (75 μM) and 0,003% SDS was added as described in (A). ALP activity was measured as described in the Material and methods. Data are an average from four independent experiments.

The same experiment was performed on cells subjected to 3 hours or 6 hours of nitrogen starvation (Fig. 5B). ALP activity was 3-fold increased after 6 hours of nitrogen deprivation without MG-132 suggesting that mitophagy was induced. However, MG-132 treatment caused only a slight enhancement of mitophagy after 3 hours or 6 hours of nitrogen starvation compared to the mitophagy of untreated cells. This suggests that mitophagy is not dependent on proteasome activity in short durations of nitrogen starvation.

Together with results published by Müller et al. (2015),^16^ our findings attest to the possibility that mitophagy can be regulated by ubiquitination/deubiquitination events and suggest that regulation mechanisms are markedly nonlinear.

### Investigation of Atg32 expression in *bre5Δ, ubp3Δ*, and *yme1Δ* mutants

When core components of the Ubp3-Bre5 deubiquitination complex^17,18^ were found to dynamically translocate from the cytosol to mitochondria upon induction of mitophagy, a role of ubiquitination in the regulation of mitophagy was suggested.^16^ Müller et al. did not characterize mitophagy proteins that were targeted by Ubp3-Bre5 complex. We therefore speculated on the involvement of Bre5 and Ubp3 proteins in the degradation of Atg32 protein in the stationary phase of growth. To test this, we expressed Atg32-V5 in *bre5Δ* and *ubp3Δ* mutants. As shown in Figure 6, the Atg32-V5 protein levels were moderately lower in *bre5Δ* and *ubp3Δ* mutants in an early exponential growth phase compared to control cells. Compared to control cells, the slightly delayed disappearance of Atg32 through the course of growth and in the stationary phase was also seen in *bre5Δ* cells. Interestingly, maximal Atg32 levels in *ubp3Δ* mutant were barely reduced compared to control cells. In parallel, we checked mitophagy induction in *bre5Δ* and *ubp3Δ* mutants expressing Idp-GFP by western blots during growth. We observed that, in the exponential growth phase, mitophagy is already induced, and during the stationary growth phase, the level of mitophagy increased very little (Fig. 7). It has been previously shown that mitophagy is disturbed in these mutants.^11,16,19^ Nevertheless, these observations suggest that expression of Atg32 and mitophagy are deregulated in these mutants and that both *Bre5Δ* and *Ubp3Δ* mutants are probably not directly involved in the Atg32 regulation by ubiquitination/deubiquitination events.

**Figure 6:**
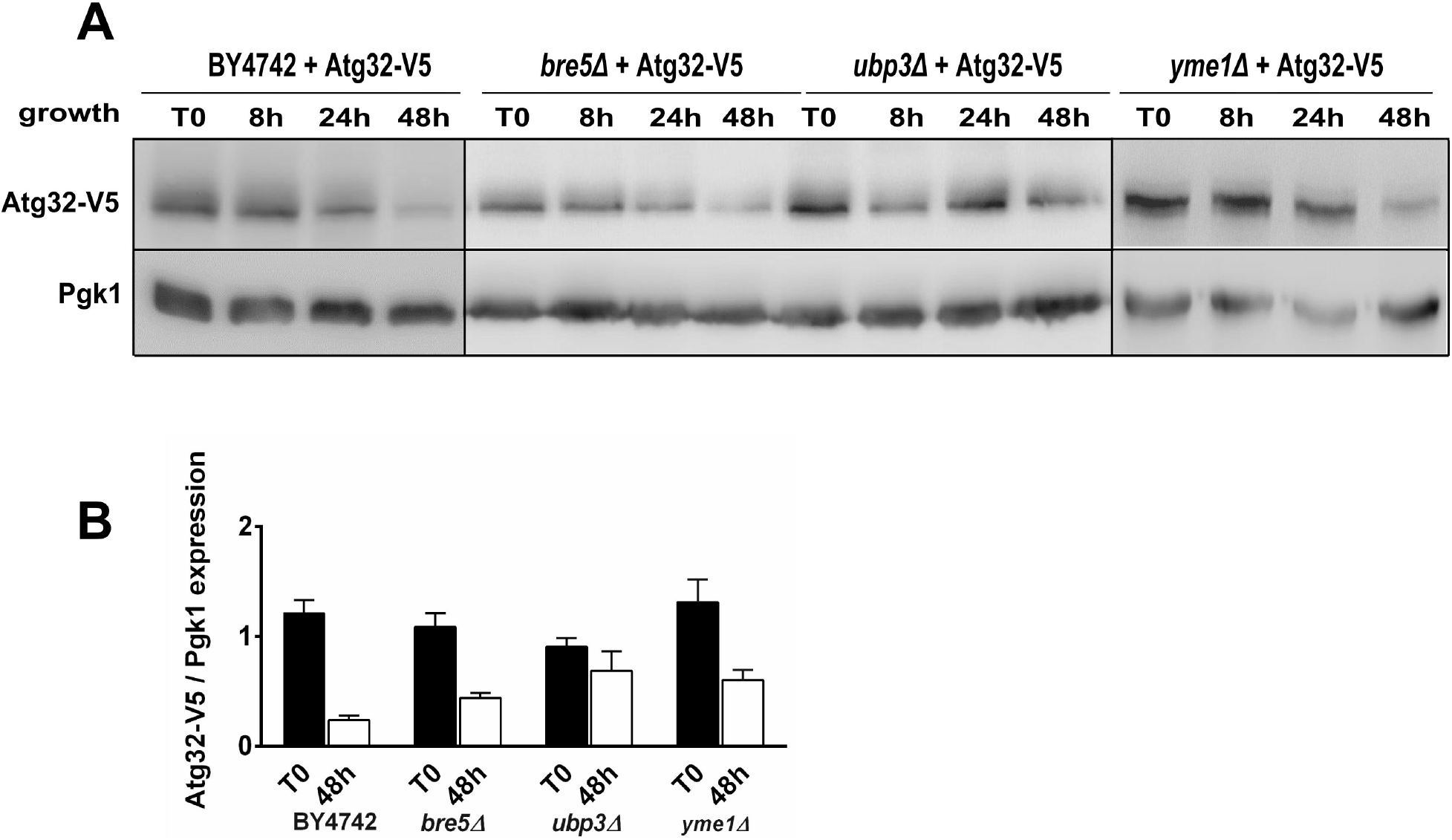
Atg32 expression is not markedly altered in deubiquitinase *bre5Δ* and *ubp3Δ* mutants strains. **(A)** Control (BY4742), *bre5Δ, ubp3Δ*, and *yme1Δ* mutant cells expressing Atg32-V5 and Idp1-GFP fusion proteins grown in a minimal synthetic medium containing lactate were harvested cells in early exponential phase of growth (T0) and then at different time points of cell growth (8h, 24h and 48h). Total protein extracts were prepared afterwards and protein samples were analyzed by western blots. Anti-V5 antibody was used to visualize Atg32-V5 recombinant protein; Pgk1, the cytosolic phosphoglycerate kinase, was used as loading control. **(B)** Atg32-V5/Pgk1 ratios were quantified at T0 and T48 for all tested strains. Three independent experiments were carried out for each condition.

It was previously shown that under certain condition the processing of Atg32 by mitochondrial i-AAA protease Yme1 acts a as an important regulatory mechanism of cellular mitophagy activity.^11^ To assess whether Yme1 may control the level of Atg32, we concurrently monitored the steady-state levels of Atg32-V5 protein (Fig. 6) and mitophagy-dependent processing of Idp-GFP (Fig. 7) in the early to stationary phase of growth in *yme1Δ* mutant under respiratory conditions. We found that the Atg32 levels diminished in the stationary phase of growth (48h) in this mutant but less than in the control cells (Fig. 6). Similarly to *bre5Δ* and *ubp3Δ* mutants, mitophagy is already induced in an early exponential growth phase in *yme1Δ* cells and has changed marginally through the growth course or in the stationary phase, respectively (Fig. 7).

**Figure 7:**
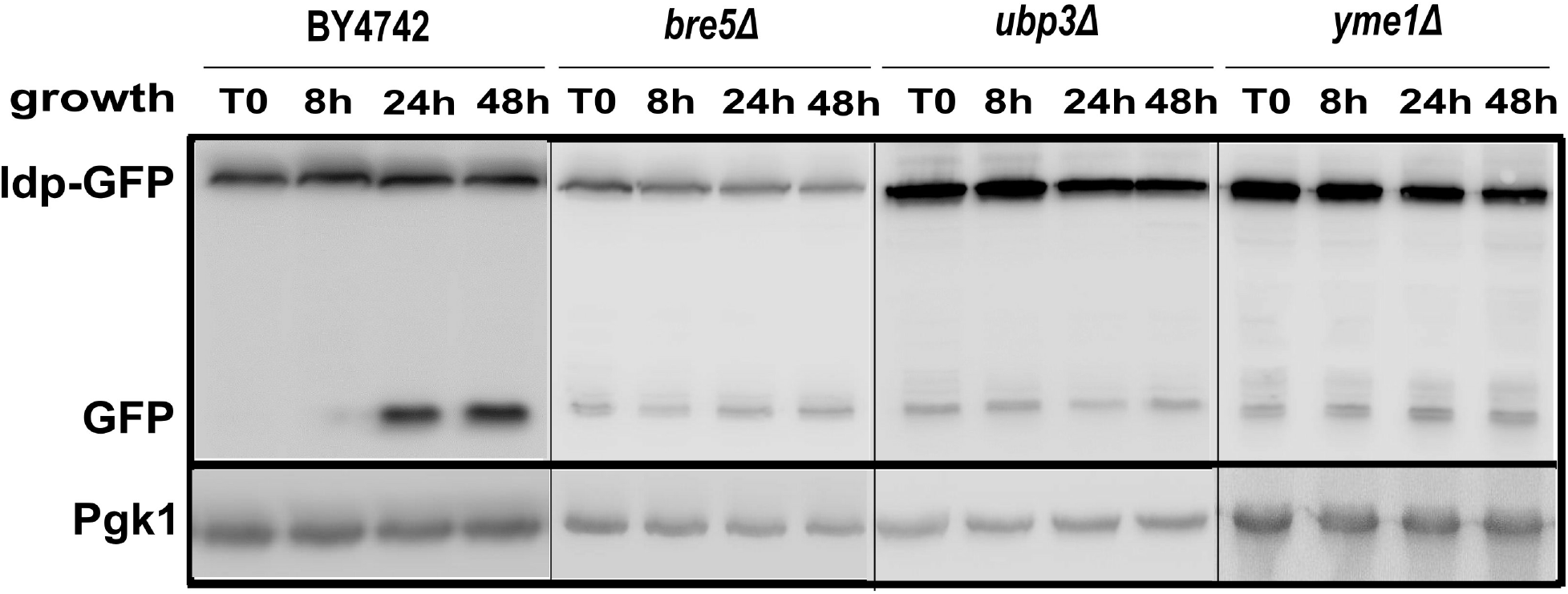
Mitophagy is altered in *bre5Δ*, *ubp3Δ*, and *yme1Δ* mutant strains. (A) Cells expressing Atg32-V5 and Idp1-GFP proteins grown in a minimal synthetic medium with lactate were harvested at different times of growth: at T0 corresponds to a mid-exponential phase of growth and then after 8-, 24- and 48 hours of cell growth. Total protein extracts from 2 × 10^7^ cells were prepared and separated on a 12,5% SDS-PAGE gel as described in the Material and methods section. Proteins were detected using antibodies against GFP (Roche) or Pgk1 (Invitrogen).

### Identification of ubiquitination site in Atg32 protein by LC-MS/MS analysis

To confirm our hypothesis that levels and activity of Atg32 protein could be also regulated by ubiquitination, we purified Atg32 protein from *atg32Δ* mutant cells expressing Atg32-V5-6xHIS harvested in the stationary growth phase in presence of MG-132. Cell lysate prepared as described in the Material and methods section, was loaded on an affinity Ni-NTA column. After elution and analysis on SDS-PAGE, two bands were detected by Coomassie blue coloration (Fig. 8A). The presence of Atg32 in band 1 was confirmed by detection with antibodies directed against histidine and ubiquitin (Fig. 8B). Nevertheless, it should be noticed that the protein did not migrate at its expected mass of 58.9 kD, but was detected in an oligomer with an high mass (larger than 170 kD) despite the presence of detergent. To ascertain Atg32 ubiquitination, the band 1 containing Atg32 protein was submitted to a classical proteomics workflow as described in the Material and Methods section including an on-gel proteolysis step and an analysis by LC-MS/MS of peptides. Specifically, trypsin digestion of ubiquitinated proteins cleaves off all but the two C-terminal glycine residues of ubiquitin from the modified protein. These two C-terminal glycine (GG) residues remain linked to the epsilon amino group of the modified lysine residue in the tryptic peptide derived from digestion of the substrate protein. The presence of the GG on the side chain of that Lys prevents cleavage by trypsin at that site, resulting in an internal modified Lys residue in a formerly ubiquitinated peptide.

**Figure 8:**
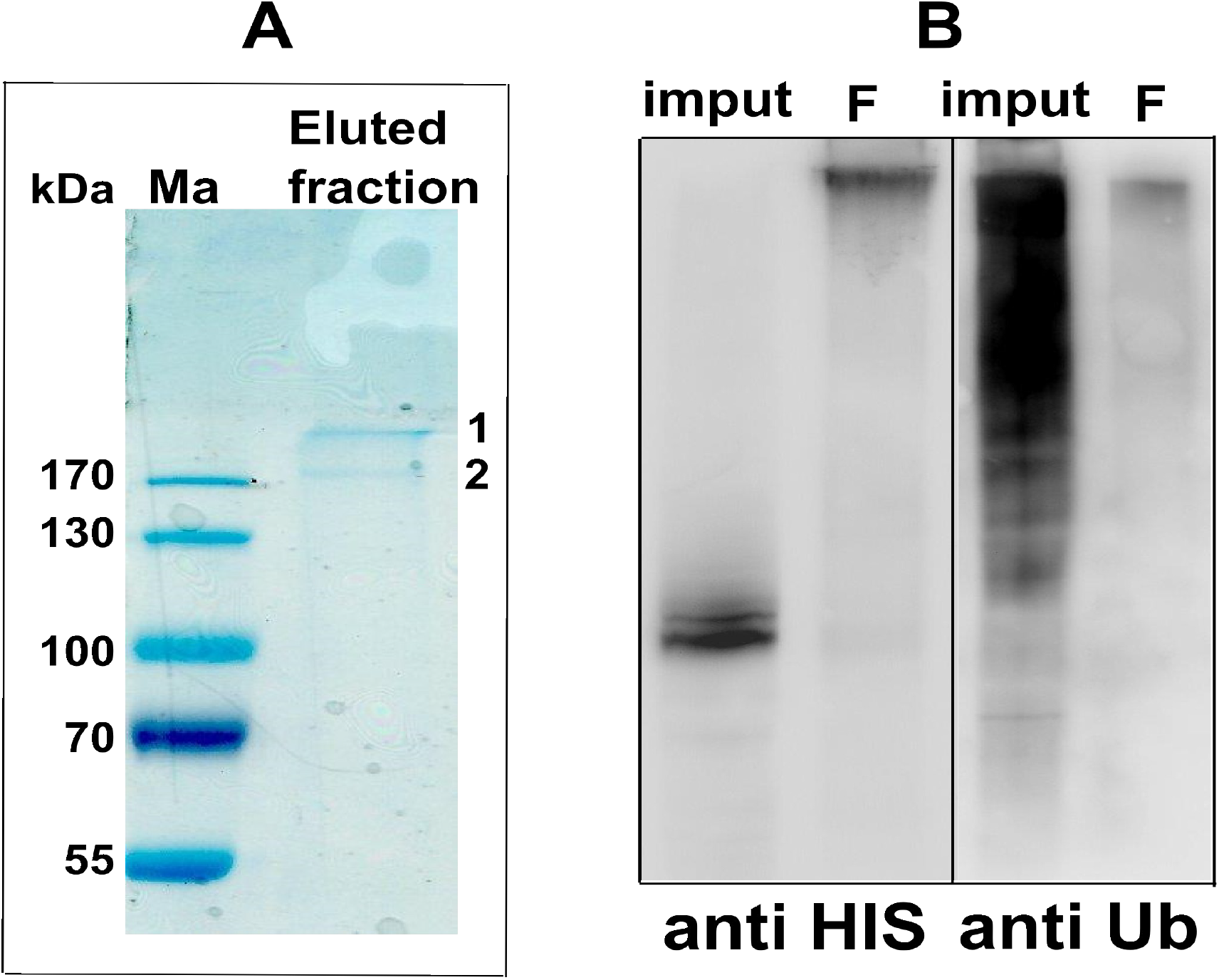
Atg32 purification and mass spectrometry analysis. (A) Cells expressing Atg32-V5-6HIS were grown until stationary phase (T48) in presence of MG-132. Cells were lysed as described in the Material and methods section and the lysate was loaded on a Ni-NTA column. After last elution, collected fractions were pooled and loaded on a 11% SDS-PAGE. After migration, the gel was colored with colloidal blue. (B) The fraction was loaded on a 11% SDS-PAGE gel and revealed by antibodies against Histidine or Ubiquitin, respectively. The band 1 was excised and analysed by mass spectrometry.

Atg32 was unambiguously identified on the basis of 21 unique peptides leading to 55.6% sequence coverage (Table 1). One of these peptides was unambiguously shown to bear the typical GG tag (+144 Da) remaining after tryptic proteolysis of ubiquitination of Lysines using two distinct search engines (SEQUEST and MASCOT) and manual validation (visual inspection of the related MS/MS spectrum). The simultaneous detection into the MS/MS spectrum of 20 b fragments and 22 y fragments (over 22 possible ones of each) ascertains the presence of a typical GG tag on the Lysine 282 and confirms at least one ubiquitination site into Atg32.

## Discussion

### Regulation of mitophagy by ubiquitination

Atg32 executes the key role of a receptor for multiple mitophagy-inducing pathways and control/govern the final turnover of mitochondria. Therefore, a thorough regulation of its activity is expected. Besides regulation of its gene expression, posttranslational modification seems to control Atg32-mediated mitophagy. However, the pathways perceiving mitochondrial damage/impairment and Atg32 activation are not fully understood.

In this work, we were interested in the relationship between mitophagy, Atg32, and the proteasome. Atg32-V5 protein was expressed in the mid-exponential phase of growth, and disappeared progressively with growth, and was almost entirely missing in the stationary phase of growth and during long durations of nitrogen starvation. Several groups already showed that Atg32 protein was expressed during growth in respiratory conditions and disappeared in the stationary phase of growth.^4,11^ Wang et al. (2013) demonstrated that Atg32 is processed by Yme1 protease, resulting in Atg32 C-terminus cleavage, and this processed form is required for mitophagy.^11^ We tagged Atg32 protein with a V5 tag in C-terminus, and this recombinant protein may be subjected to Yme1-dependent processing, resulting in the loss of the V5 tag, upon mitophagy induction. However, in Wang et al.’s study, Atg32 processing was very weak and slow in cells grown in the presence of lactate as carbon source; about 5% of Atg32 was cleaved by Yme1 in both nitrogen starvation and in the stationary phase of growth. Moreover, in the same article the authors showed the disappearance of Atg32 tagged in N-terminus with a TAP tag in the stationary phase of growth, and we observed the same behavior with HA-Atg32. Therefore, our results concerning the disappearance of Atg32-V5 in the stationary phase of growth are in line with the published data, and Yme1 probably plays a minor role in the disappearance of Atg32-V5 recombinant protein in this phase.

Considering the important role of the proteasome in the regulation of numerous proteins of the outer mitochondrial membrane,^20^ we assessed whether the proteasome is also involved in Atg32 degradation. Indeed, blocking the proteolytic activity of proteasome by MG-132 allowed us to prevent a decline in the Atg32 levels through the course of growth and in the stationary phase and to stabilize the modified form of Atg32 (Fig. 3). Moreover, in the proteasome *pre2-2* mutant strain, the Atg32 levels decreased much less compared to control cells (Fig. 4) These results indicate that the Atg32 protein levels are dependent on proteasome activity.

It was previously demonstrated that overexpression of Atg32 stimulates mitophagy and it has long been known that the absence of Atg32 protein causes mitophagy impairment in yeast.^3,4^ These published data suggest that mitophagy is dependent on the amount Atg32 protein present. Our results are in line with this observation. Mitophagy was stimulated upon MG-132 treatment in the stationary phase of growth and we demonstrated that Atg32-V5 protein levels were more elevated in this condition than in untreated cells. The proteasome may regulate Atg32 protein amount so that yeast cells can control mitophagy level.

Altogether, our data indicate that, as in mammalian cells, yeast mitophagy can be regulated by ubiquitination-deubiquitination events, and an interplay exists between mitochondria degradation by autophagy and the proteasome. It was shown that outer mitochondrial membrane ubiquitin ligases, such as mammalian MULAN, MARCHV/MITOL, and yeast Mdm30, can ubiquitinate proteins involved in mitochondrial fusion and fission, targeting them for proteasome degradation, thus affecting mitochondrial dynamics.^21,22,23^ These ligases ensure outer mitochondrial membrane quality control. Atg32-V5 expression seems to be dependent on proteasome activity, suggesting that Atg32 is a substrate of this degradation machinery and can be ubiquitinated.

The deubiquitination enzymes (DUBs) are highly diverse functionally, reflecting both their subcellular localization and their inherent substrate specificities. An important function of the DUBs is to recycle ubiquitin by recovering it from ubiquitin–protein conjugates before the target protein is degraded. Defects in this process give rise to reduced ubiquitin levels and pleiotropic stress sensitivities. The main DUBs responsible for recovering ubiquitin from conjugates that are en route to being degraded are Ubp6, Rpn11 and Doa4.^24,25,26,27,28,29,30^ To our knowledge, it is not known what is the contribution of Ubp3 protein to cellular deubiquitylation. But is known that it cleaves ubiquitin fusions but not polyubiquitin.

Moreover, we do not know if Atg32 is mono- or poly-Ub. However, we can assume that Bre5-Ubp3 complex participates in reducing ubiquitin level to some extent in the cells. Therefore, we can speculate that lowered availability of free Ub for modification of protein target in *ubp3Δ* could result in accumulation of protein target, which was not modified with Ub and thus degraded by proteasome. From this perspective, our result (Figure 6: the Atg32 level is increased in *ubp3Δ* mutant compared with control cells in stationary growth phase) is not so surprising.

Ubiquitination of mitochondrial proteins can play two different roles: (i) ubiquitination of proteins allows their turnover by proteasomal degradation and, consequently their quality control; and (ii) ubiquitination of proteins localized on the outer mitochondrial membrane is a signal to trigger mitophagy. In the case of Atg32 mitophagy receptor, different post-translational modifications take place to modulate its activity. The fact that activity of Atg32 may be regulated by the proteasome is another example of the existence of a dialogue between these two degradative processes.

### Why is Atg32 regulated by multiple post-translational modifications?

During mitophagy induction Atg32 is activated by a posttranslational modification. There is still the question of why Atg32 protein is subjected to several different post-translational modifications, such as phosphorylations, ubiquitination, or other unknown modifications, and what the roles of these translational modifications are. Are these modifications dependent on mitophagy induction conditions (e.g., nutrient starvation, stationary phase of growth, or rapamycin treatments)? From published data, phosphorylation of Atg32 at serine 114 and serine 119 by a serine/threonine protein kinase Casein kinase 2 (CK2) seems to be required for the interaction with the adaptor protein Atg11 and targeting mitochondria for the degradation into vacuole.^7^ Thus, vacuolar degradation is responsible for Atg32 stability bearing this kind of modification (Fig. 3A, ± PMSF). What is the fate of Atg32 on mitochondria that have not been selected for mitophagy? We assumed that Atg32 could be eliminated by a separate pathway of degradation in its unmodified form or after acquiring another type of modification. In fact, our results suggest that Atg32 could also be ubiquitylated, and mitophagy can be regulated by ubiquitination/deubiquitination events. We were able to confirm ubiquitination of Atg32 protein at least on lysine in position 282 by LC-MS/MS analysis (Fig. 8, Fig. S5 and Table 1). At this point, it would be interesting to examine if Atg32 ubiquitination is dependent on Atg32 phosphorylation. We can hypothesize that ubiquitination will be useful to regulate Atg32’s own expression and to control mitophagy level. This would prevent excessive mitochondria degradation and ensure a fine-tuned mitochondria turnover (Fig. 9). In mammalian cells, such a situation was already observed in the case of FUNDC1 receptor, which is principally involved in mitophagy during hypoxia. This receptor is regulated by both phosphorylation and ubiquitination.^31,32,33,34,35^ The mitochondrial PGAM5 phosphatase interacts with and dephosphorylates FUNDC1 serine 13 (Ser-13) residue upon hypoxia or carbonyl cyanide p-trifluoromethoxyphenylhydrazone (FCCP) treatment. Dephosphorylation of FUNDC1 catalyzed by PGAM5 enhances its interaction with LC3. CK2 phosphorylates FUNDC1 to reverse the effect of PGAM5 in mitophagy activation. Indeed, the mitochondrial E3 ligase MARCH5 plays a role in regulating hypoxia-induced mitophagy by ubiquitinating and degrading FUNDC1 to limit excessive mitochondria degradation (Fig. 7).^32^ We believe that all these open questions are important in the study of mitophagy. Obviously, additional experiments are needed to further investigate how cells control the level and activity of Atg32 to be able to understand the mechanisms by which cells control selective degradation of mitochondria, and the physiological significance of mitophagy. In addition, an interesting question is whether Atg32 is also involved in other cellular processes. Future studies in yeast will address these important questions.

**Figure 9:**
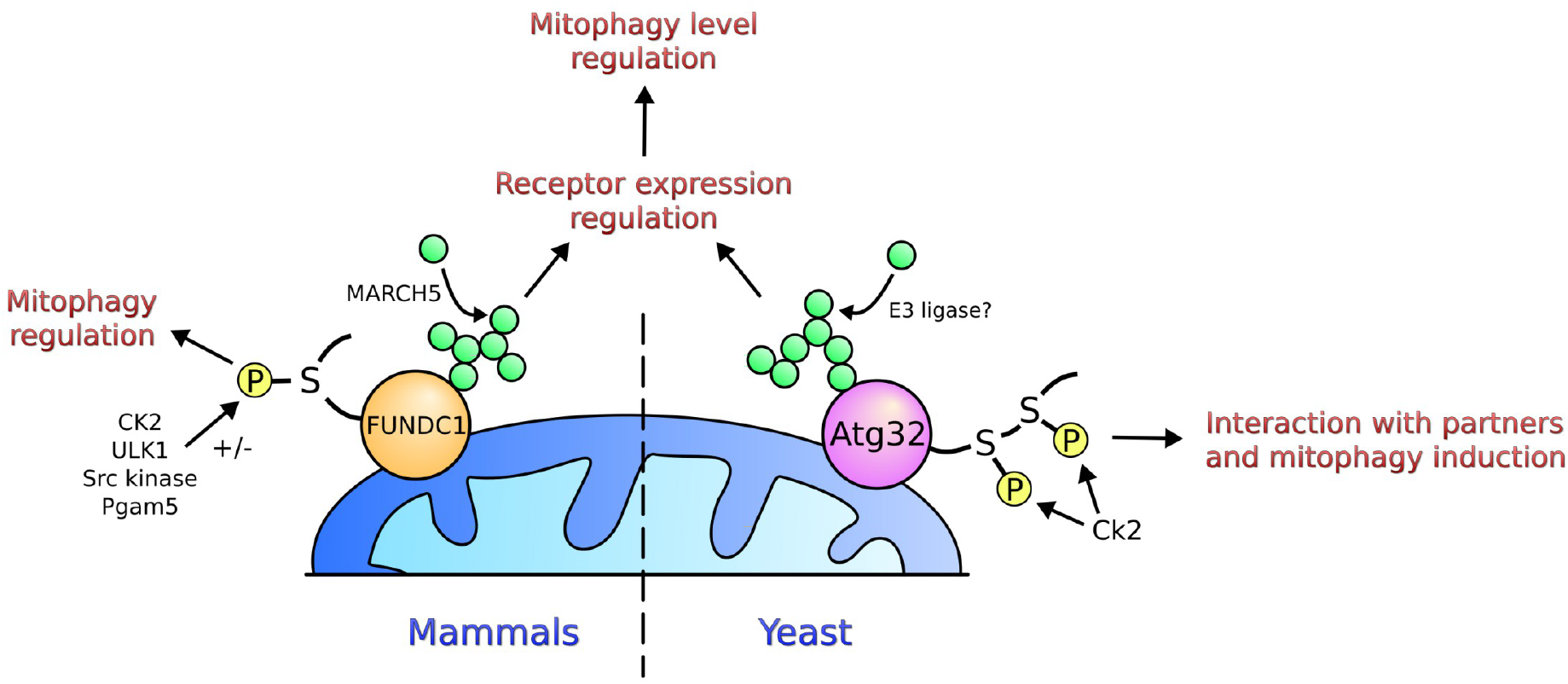
Post-translational modifications of yeast Atg32 and mammalian FUNDC1 mitophagy receptors. The mitochondrial PGAM5 phosphatase interacts with and dephosphorylates FUNDC1 at serine 13 (Ser-13) upon hypoxia or FCCP treatment. Dephosphorylation of FUNDC1 catalyzed by PGAM5 enhances its interaction with LC3. CK2 phosphorylates FUNDC1 to reverse the effect of PGAM5 in mitophagy activation. ULK1 also interacts with FUNDC1, phosphorylating it at serine 17, enhancing FUNDC1 binding to LC3. In yeast, Atg32 is phosphorylated on serines 114 and 119 by Ck2 kinase; this modifications are crucial for the initiation of mitophagy. Our results indicate that Atg32 is also regulated by ubiquitination.

## Material and methods

### Yeast strains, plasmids and growth conditions

All yeast strains used in this study are listed in the supplementary Table I and derived from BY4742 (Euroscarf bank). Yeast cells were grown aerobically at 28°C in minimal medium (0.175% yeast nitrogen base without amino acids and ammonium sulfate, 0.5% ammonium sulfate, 0.1% potassium phosphate, 0.2% Drop-Mix, 0.01% of auxotrophic amino acids and nucleotide, pH 5.5), supplemented with 2% lactate as a carbon source. Cell growth was followed by optical density at 600 nm. For starvation experiments, cells were harvested at the early exponential phase of growth, washed three times with water and incubated in nitrogen starvation medium (0.175% yeast nitrogen base without amino acids and ammonium sulfate, and 2% lactate, pH 5.5). Mitophagy was studied in cells subjected to nitrogen starvation for 3, 6 or 24 hours or during different phases of growth and expressing the mitochondrial matrix Idp1-GFP protein. The plasmid p*IDP1-GFP* was a gift from Dr. Abeliovich (Hebrew University of Jerusalem, Israel). The plasmid expressing Atg32-V5-6xHIS from ATG32 promoter and HA-Atg32 with copper promoter were gifts from Axel Athané (IBGC, Bordeaux) and Dr. Dan Klionsky (Michigan University, USA) respectively. The promoter of ATG32 gene was amplified by polymerase chain reaction (PCR) using forward primer (5’-GTGATGTATCCACAGGGAATTCCGCTC-3’) and reverse primer (5’-CTTTTAGATGAGGATCCTTTACCT-3’) and cloned into BamH1-digested YEp357 plasmid kindly provided by Dr Pinson (IBGC, Bordeaux) to obtain the pPROM-*ATG32-β galactosidase* plasmid.

### Preparation of protein extracts and western blots

For preparation of total protein extract, 2×10^7^ cells were harvested by centrifugation, washed with water and resuspended in 450 μl of water and 50 μl of lysis buffer (1.85 M NaOH, 3.5% β-mercaptoethanol). After 10 min on ice, 50 μl of trichloroacetic acid 3 M was added followed by another incubation of 10 min on ice. Proteins were pelleted by centrifugation, 8 min at 13 000xg, washed with acetone and resuspended in 20 μl of 5% SDS and 20 μl of loading buffer (2% β-mercaptoethanol, 2% SDS, 0.1 M Tris-HCl, pH 8.8, 20% glycerol, 0.02% bromophenol blue). Samples were boiled 5 min and 50 μg of proteins were separated by electrophoresis on 12.5% SDS-PAGE and subjected to immunoanalysis with either anti-GFP antibody (Roche), anti-Pgk1 antibody (Invitrogen), anti-porin (Invitrogen), anti-Dpms (Invitrogen), anti HA (Roche), anti-V5 (Invitrogen) and anti-ubiquitin (Calbiochem). Detection was performed with ECL^+^ reagent (Perkin Elmer) and western blot quantifications were done using ImageJ Software (NIH). Results of quantification were expressed as the mean ± SEM. For preparation of cell lysates, 5×10^7^ cells were broken with glass beads in buffer containing 0,6 M sorbitol, 20 mM MES pH 6 plus protease inhibitor cocktail (Roche); lysates were centrifuged 10 min at 800xg. The supernatants were loaded on 20-55% OptiPrep™ density gradients in buffer with 0,6M sorbitol 20 mM MES pH6, 5 mM EDTA, pH6 plus protease inhibitor cocktail (Roche). Fractions were collected, precipitated with TCA and pellets were resuspended in 20 μl of 5% SDS and 20 μl of loading buffer (2% β-mercaptoethanol, 2% SDS, 0.1 M Tris-HCl, pH 8.8, 20% glycerol, 0.02% bromophenol blue). Samples were boiled 5 min and 50 μg of proteins were separated by electrophoresis on 12.5% SDS-PAGE and subjected to immunoanalysis.

### ALP activity

Alkaline phosphatase activity was performed on *pho8Δ* strain, expressing the mitochondria-targeted truncated version of Pho8, mtPho8Δ60, during the exponential (T0) and the stationary phase (T48h) of growth or after 3 and 6 hours of nitrogen starvation. For each point, five OD_600nm_ of cells expressing mitochondria-targeted Pho8Δ60 (mtPho8Δ60) were harvested and lysed with glass beads into lysis buffer (20 mM PIPES, 0.5% Triton X-100 0.5%, 50 mM KCl, 100 mM potassium acetate, 10 mM MgCl_2_, 10 μM ZnSO_4_, 2 mM PMSF). After centrifugation, 20 μl of supernatant were put with 80 μl of water and 400 μl of activity buffer (250 mM Tris HCl pH 8, 0.4% Triton X-100, 10 mM MgCl_2_, 10 μM ZnSO_4_, 125 mM p-nitrophenyl-phosphate) during 20 minutes at 30°C. The reaction was stopped by the addition of 500 μl of 1 M glycine pH 11. Activity was then measured by optical density at 400 nm. Protein concentration was measured with the Lowry method. ALP activities were expressed as arbitrary fluorescence units/minute/mg proteins (AU/min/mg).

### β-galactosidase activity

BY4742 wild-type cells, expressing β-galactosidase protein under control of *ATG32* promoter were grown in a lactate-containing medium and harvested at different time points: midexponential phase of growth (T0), late exponential phase of growth (8h), early stationary phase of growth (24h) and late stationary phase of growth (48h). For proteasome inhibition, 75μM MG-132 + 0,003% SDSwere added at 8h time point. To measure β-galactosidase activity, five OD_600nm_ (corresponding to 5.0×10^7^ cells) were harvested and broken with glass beads in lysis buffer (100mM Tris-HCL pH 8, 1mM DTT, 20% glycerol, 2mM PMSF) during 4min at 20 hertz using Mixer Mill MM400 (Retsch). Then, 20μl of cell lysate were used to measure β-galactosidase activity by absorbance at 420nm in 60mM Na_2_HPO_4_, 40mM NH_2_PO_4_, 10mM KCl, 1mM MgSO_4_, 50mM β-mercaptoethanol, pH 7 and with 0.8mg ortho-Nitrophenyl-β-galactoside (ONPG). Protein concentrations were measured using Lowry’s method and activities were normalized to T0.

### Atg32 purification

To purify Atg32 protein, 20 OD cells expressing Atg32-V5-6xHIS and grown until stationary phase in presence of 75 μM MG-132 were collected and lysates were prepared as described in Becuwe et al.^36^ Cells were precipitated with TCA to a final concentration of 10% overnight at 4°C, and lysed with glass beads for 20 min at 4°C. After centrifugation, the lysate was collected and centrifuged at 16000xg for 10 min. The pellet was resuspended with 30 μl of 1 M non buffered Tris and 200 μl of guanidium buffer (6 M GuHCl, 20 mM Tris-HCl, pH 8, 100 mM K_2_HPO_4_, 10 mM imidazole, 100 mM NaCl, and 0.1% Triton X-100) and incubated for 1 h at room temperature on a rotating platform. After centrifugation (16,000xg for 10 min at room temperature), the lysate was loaded on a HisTrap FF Crude column (GE healthcare Life Sciences). The column was then washed with a speed of 1 ml/min with guanidium buffer, then with wash buffer 1 (20 mM Tris-HCl, pH 8.0, 100 mM K_2_HPO_4_, 20 mM imidazole, 100 mM NaCl, and 0.1% Triton X-100) and then with wash buffer 2 (20 mM Tris-HCl, pH 8.0, 100 mM K_2_HPO_4_, 10 mM imidazole, 1 M NaCl, and 0.1% Triton X-100). His6-tagged proteins were finally eluted with elution buffer (50 mM Tris-HCl, pH 8.0, and 250 mM imidazole). Fractions absorbing at 254nm were pooled and analyzed by electrophoresis on 11% SDS-PAGE and submitted to blue colloidal staining and immuno-analysis.

### Sample preparation and protein digestion for MS analysis

After colloidal blue staining, the band corresponding to Atg32 was cut out from the SDS-PAGE gel and subsequently cut in 1 mm × 1 mm gel pieces. Gel pieces were destained in 25 mM ammonium bicarbonate 50% acetonitrile (ACN), rinsed twice in ultrapure water and shrunk in ACN for 10 min. After ACN removal, gel pieces were dried at room temperature, covered with the trypsin solution (10 ng/μl in 50 mM NH_4_HCO_3_), rehydrated at 4 °C for 10 min, and finally incubated overnight at 37 °C. Spots were then incubated for 15 min in 50 mM NH_4_HCO_3_ at room temperature with rotary shaking. The supernatant was collected, and an H_2_O/ACN/HCOOH (47.5:47.5:5) extraction solution was added onto gel slices for 15 min. The extraction step was repeated twice. Supernatants were pooled and concentrated in a vacuum centrifuge to a final volume of 100 μL. Digests were finally acidified by addition of 2.4 μL of formic acid (5%, v/v) and stored at −20 °C.

### nLC-MS/MS analysis

Peptide mixture was analyzed on a Ultimate 3000 nanoLC system (Dionex, Amsterdam, The Netherlands) coupled to a Electrospray Orbitrap Fusion™ Lumos™ Tribrid™ Mass Spectrometer (Thermo Fisher Scientific, San Jose, CA). Ten microliters of peptide digests were loaded onto a 300-μm-inner diameter × 5-mm C_18_ PepMap™ trap column (LC Packings) at a flow rate of 10 μL/min. The peptides were eluted from the trap column onto an analytical 75-mm id × 50-cm C18 Pep-Map column (LC Packings) with a 4–40% linear gradient of solvent B in 45 min (solvent A was 0.1% formic acid and solvent B was 0.1% formic acid in 80% ACN). The separation flow rate was set at 300 nL/min. The mass spectrometer operated in positive ion mode at a 1.8-kV needle voltage. Data were acquired using Xcalibur 4.1 software in a data-dependent mode. MS scans (*m/z* 375-1500) were recorded at a resolution of R = 120 000 (at m/z 200) and an AGC target of 4 × 10^5^ ions collected within 50 ms. Dynamic exclusion was set to 60 s and top speed fragmentation in HCD mode was performed over a 3 s cycle. MS/MS scans with a target value of 3 × 10^3^ ions were collected in the ion trap with a maximum fill time of 300 ms. Additionally, only +2 to +7 charged ions were selected for fragmentation. Others settings were as follows: no sheath nor auxiliary gas flow, heated capillary temperature, 275 °C; normalized HCD collision energy of 30% and an isolation width of 1.6 m/z. Monoisotopic precursor selection (MIPS) was set to Peptide and an intensity threshold was set to 5 × 10^3^.

## Acknowledgments and fundings

The work in the laboratories of the authors was supported by grants from the Slovak APVV agency (SK-FR-2015-0005) (to IBK), the CNRS, the University of Bordeaux and the Doctoral International Program from IDEX of Bordeaux supported by Agence Nationale pour la Recherche (ANR-10-IDEX-03-02) (to NC and PV). France/Slovakia collaboration was supported by Campus France (Stefanik n° 35809VC) (to NC and IBK). We thank Drs Sagot, Klionsky, Pinson and Abeliovich, and Mr Athané for the gift of strains and material and Drs Chaignepain and Claverol (CGFB, Bordeaux) for helpful discussions.

