## Supplementary data for "Ubiquitination of the yeast receptor Atg32 modulates mitophagy"

**Figure S1: Expression of HA-Atg32 protein. (A)** *atg32Δ* mutant cells grown in lactate-containing medium and expressing HA-Atg32 recombinant protein were harvested in a mid-exponential phase of growth (T0) and after 8-, 24-, and 48 hours from T0. Total protein extracts were prepared afterwards and protein samples were analyzed by western blots. Anti-HA antibody was used to visualize HA-Atg32 recombinant protein and Pgk1, the cytosolic phosphoglycerate kinase, was used as loading control. **(B)** Atg32-V5 expression was quantified as the percentage of Atg32-V5 level of T0 (100%). Experiments were carried out three times independently.

**Figure S2: Study of Atg32-V5 expression in autophagy mutants. (A)** *atg5Δ*, *atg8Δ,* and *atg11Δ* mutant cells transformed with a plasmid expressing Atg32-V5 were grown in a minimal synthetic medium containing lactate. Cells were harvested in early an exponential phase of growth (T0) and after 8-, 24-, and 48 hours of growth. **(B)** Atg32-V5/Pgk1 ratios were quantified at T0 and T48 for all tested strains. Three independent experiments were carried out for each condition. **(C)** *atg5Δ* cells expressing Atg5-V5 were treated with MG-132 (75 µM) and 0,003% SDS at time point 8h. Total protein extracts were prepared afterwards and protein samples were analyzed by western-blots. Anti-V5 antibody was used to visualize Atg32-V5 recombinant protein and Pgk1, the cytosolic phosphoglycerate kinase, was used as loading control.

**Figure S3: The effect of MG-132 on rapamycin treatment and the Atg32 protein level in control strain (A)** *atg32Δ* cells grown in a lactate-containing medium and expressing Atg32-V5 recombinant protein were harvested at T0 and treated with 1 µg/ml rapamycin was added to the cell culture for 3, 6 and 24h +/- 75 µM MG-132 and 0,003 % SDS. Total protein extracts were prepared afterwards and protein samples were analyzed by western blots. Anti-V5 antibody was used to visualize Atg32-V5 recombinant protein; Pgk1, the cytosolic phosphoglycerate kinase, was used as loading control. **(B)** BY4742 (wild-type) cells transformed with a plasmid expressing Atg32-V5 grown in a minimal synthetic medium supplemented with 2% lactate as a carbon source were harvested in an early exponential phase of growth (T0) and after 8-, 24-, and 48 hours of cell growth. To inhibit proteasome, 75 µM MG-132 and 0,003 % SDS were added to the cell culture at 8h time point. Total protein extracts were prepared afterwards and protein samples were analyzed by western blots. Anti-V5 antibody was used to visualize Atg32-V5 recombinant protein and Pgk1, the cytosolic phosphoglycerate kinase, was used as loading control. **(C)** Atg32-V5 expression was quantified as the percentage of Atg32-V5 level of T0 (100%). Experiments were carried out three times independently.

**Figure S4: Effect of drugs treatment on growth and Atg32-V5 expression. (A)** Addition of proteasome inhibitor MG-132 (75 µM MG-132 + 0,003 % SDS) and inhibitor of vacuolar proteolysis PMSF (2 mM) do not affect growth and growth yield in *atg32Δ* mutant expressing Atg32-V5 plasmid and grown in a minimal synthetic medium containing lactate. Y-axis is represented in logarithmic scale (n = 5 for control and MG-132 ; n = 3 for PMSF). **(B)** *atg32Δ* cells grown in a lactate-containing medium and expressing Atg32-V5 recombinant protein were harvested at different time points: T0 that represents a mid-exponential phase of growth and then after 24- and 48 hours of cell growth. To inhibit proteasome, 75 µM MG-132 + 0,003 % SDS were added to the cell culture at 8 h time point (T8). To inhibit vacuolar proteolysis, 2mM PMSF was added to the cell culture at T8.. Total protein extracts were prepared afterwards and protein samples were analyzed by western blots. Anti-V5 antibody was used to visualize Atg32-V5 recombinant protein; Pgk1, the cytosolic phosphoglycerate kinase, was used as loading control.

**Figure S5: Mass spectrometry analysis.** (A) SEQUEST spectra, (B) MASCOT spectra, and (C) Protein coverage.

### Figure S1

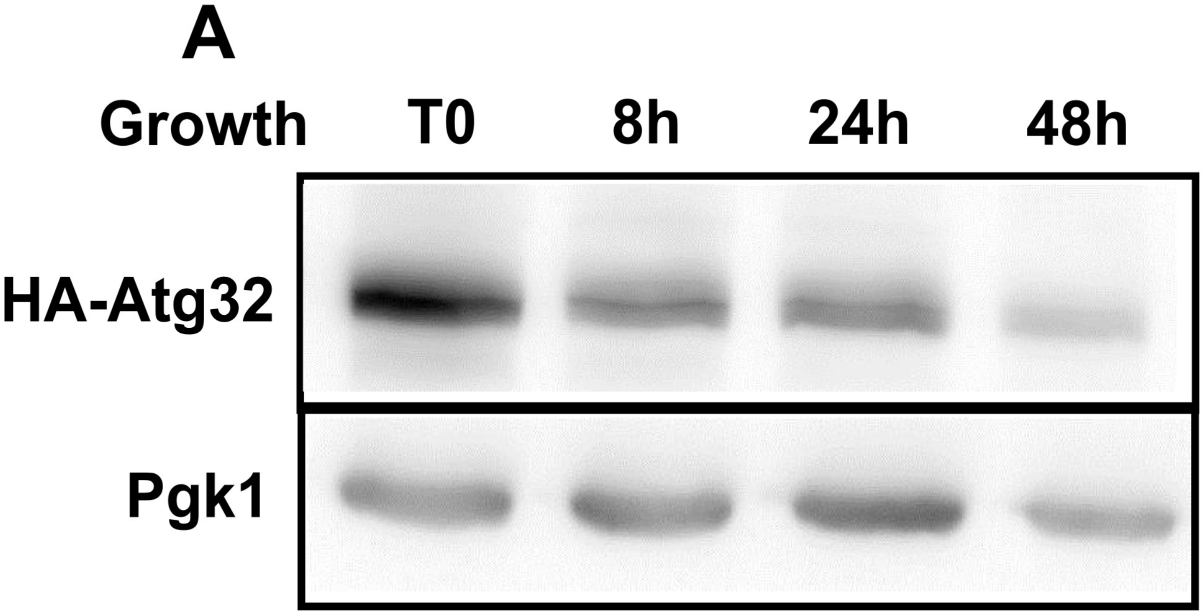
A
Growth
T0
8h
24h
48h
HA-Atg32
Pgk1

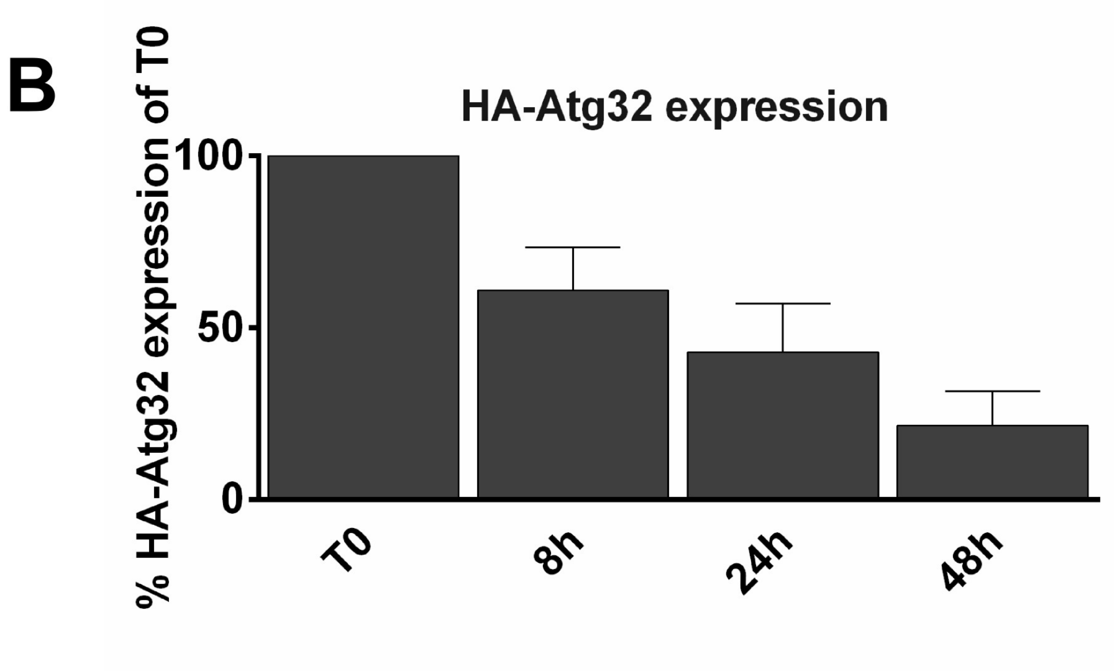
B
HA-Atg32 expression
% HA-Atg32 expression of T0
100
50
0
T0
8h
24h
48h

### Figure S2

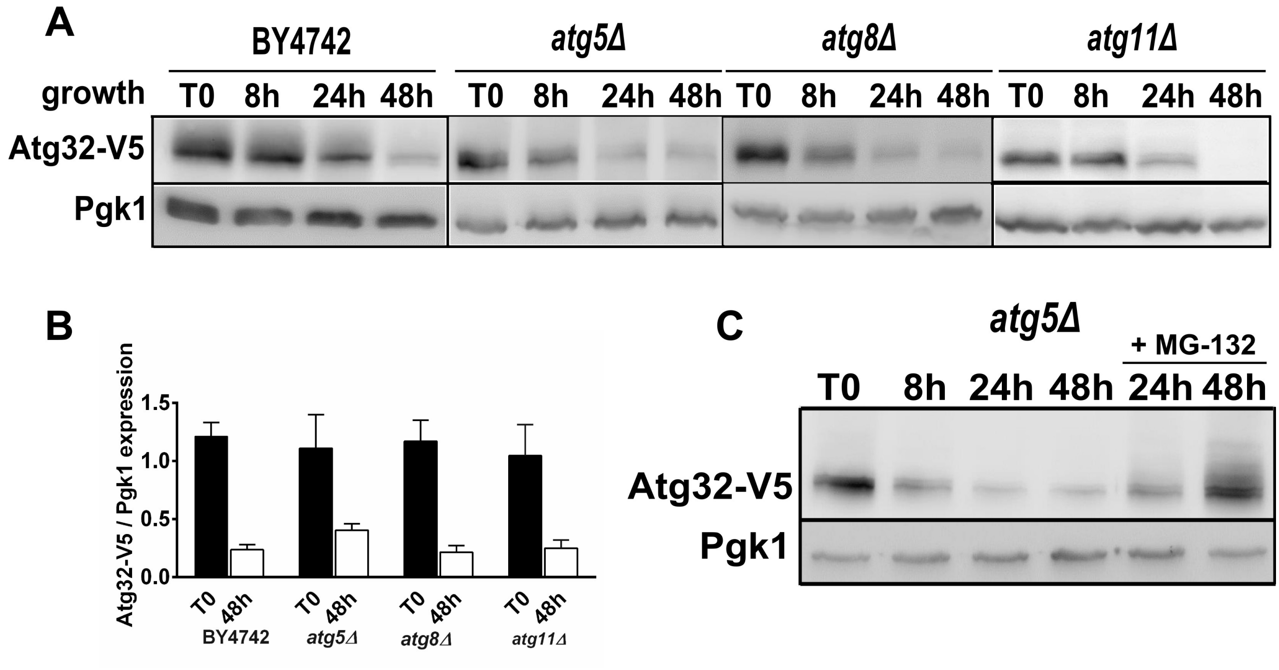
A
BY4742
atg5Δ
atg8Δ
atg11Δ
growth
T0 8h 24h 48h T0 8h 24h 48h T0 8h 24h 48h T0 8h 24h 48h
Atg32-V5
Pgk1
B
Atg32-V5 / Pgk1 expression
1.5
1.0
0.5
0.0
T0 48h
BY4742
atg5Δ
atg8Δ
atg11Δ
C
atg5Δ
+ MG-132
T0 8h 24h 48h 24h 48h
Atg32-V5
Pgk1

### Figure S3

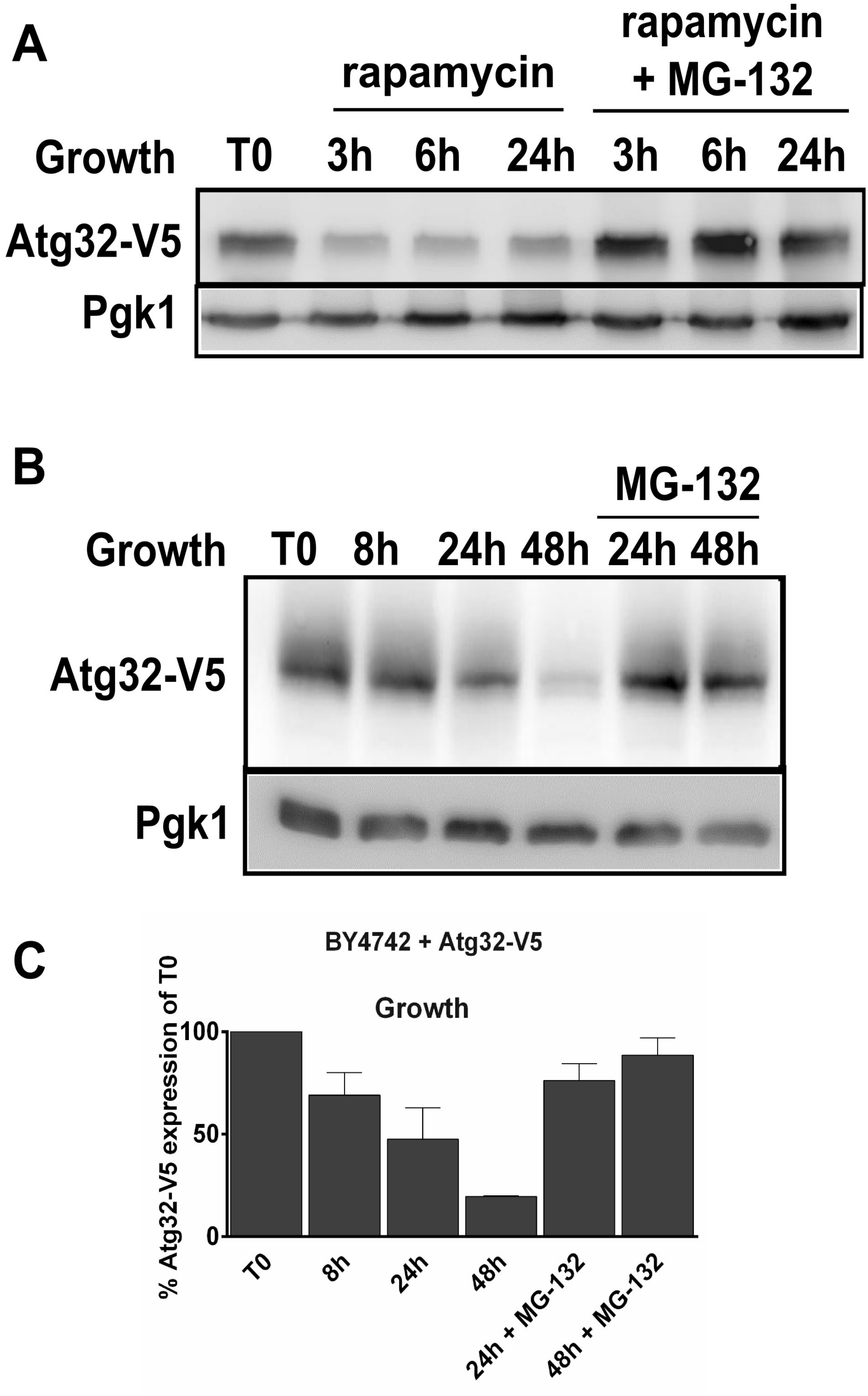
A
rapamycin
rapamycin
+ MG-132
Growth
T0
3h
6h
24h
3h
6h
24h
Atg32-V5
Pgk1
B
MG-132
Growth
T0
8h
24h
48h
24h
48h
Atg32-V5
Pgk1
C
BY4742 + Atg32-V5
Growth
% Atg32-V5 expression of T0
100
50
0
T0
8h
24h
48h
24h + MG-132
48h + MG-132

### Figure S4

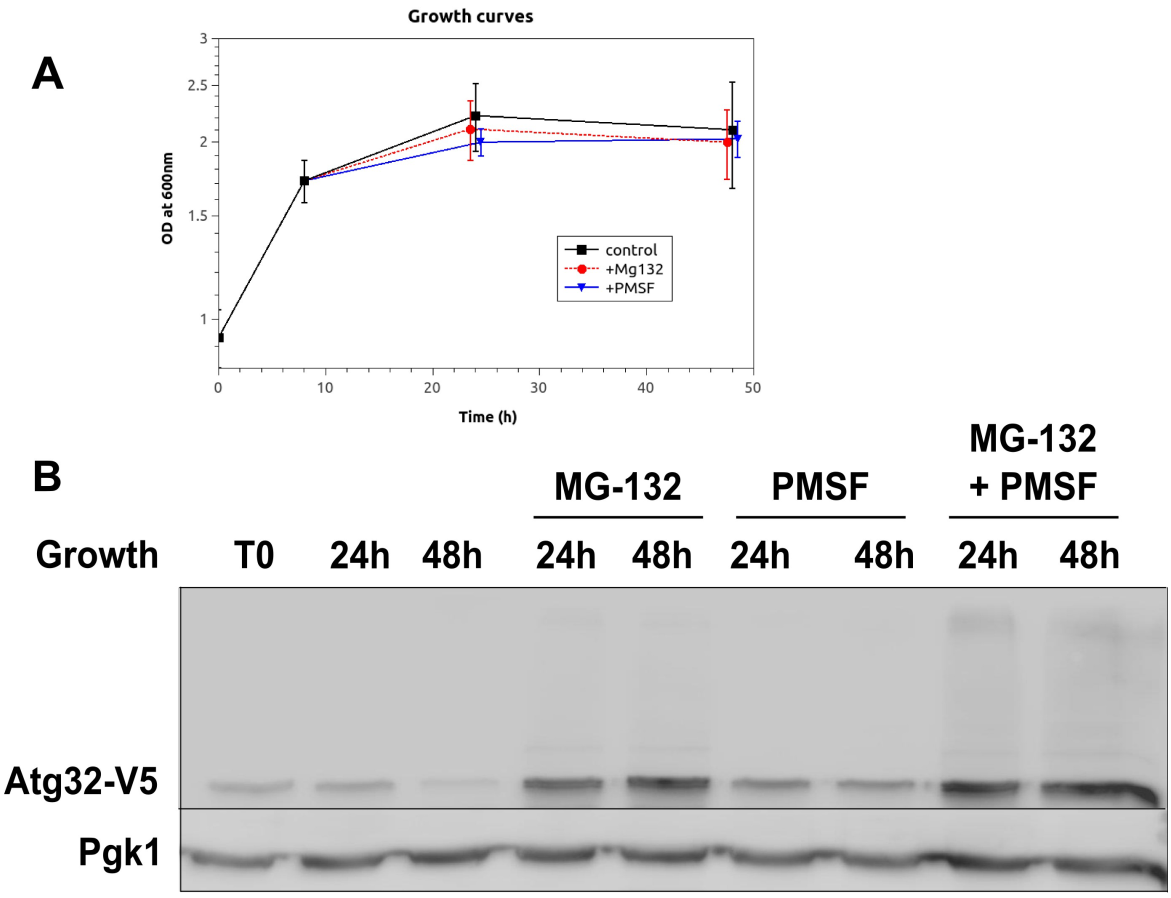

A
Growth curves
OD at 600nm
3
2.5
2
1.5
1
0
10
20
30
40
50
Time (h)
control
+Mg132
+PMSF
B
MG-132
PMSF
MG-132
+ PMSF
Growth
T0
24h
48h
24h
48h
24h
48h
24h
48h
Atg32-V5
Pgk1

### Figure S5

#### Peptide SEQUEST

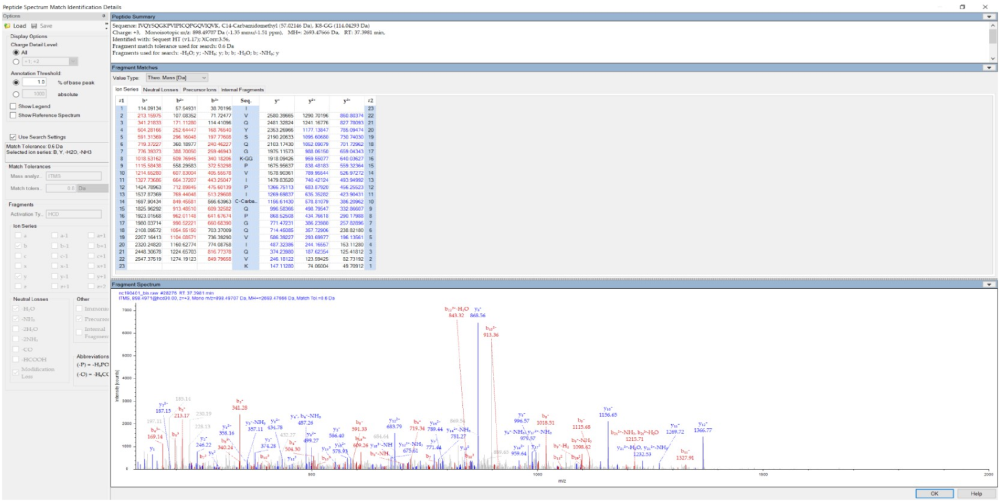

#### Peptide MASCOT

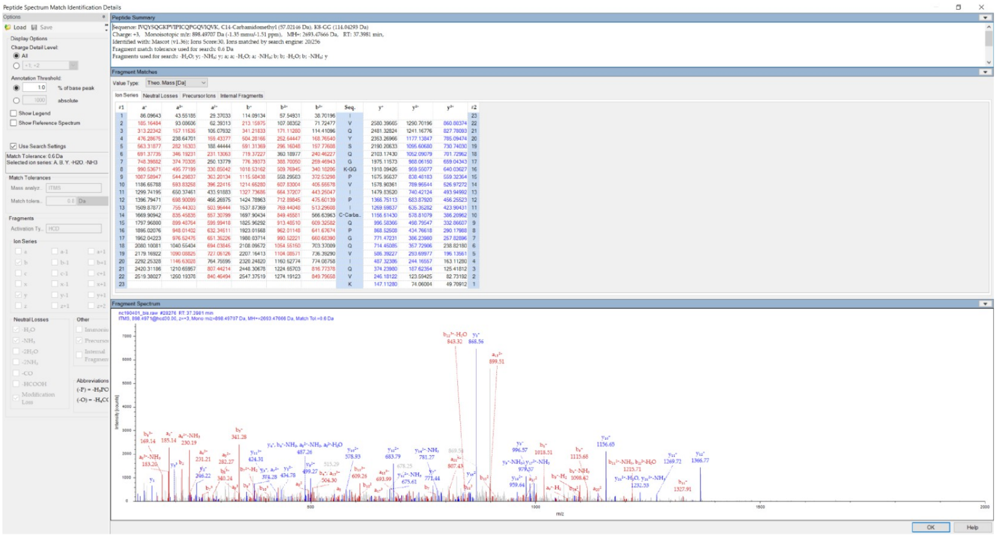

#### Protein coverage

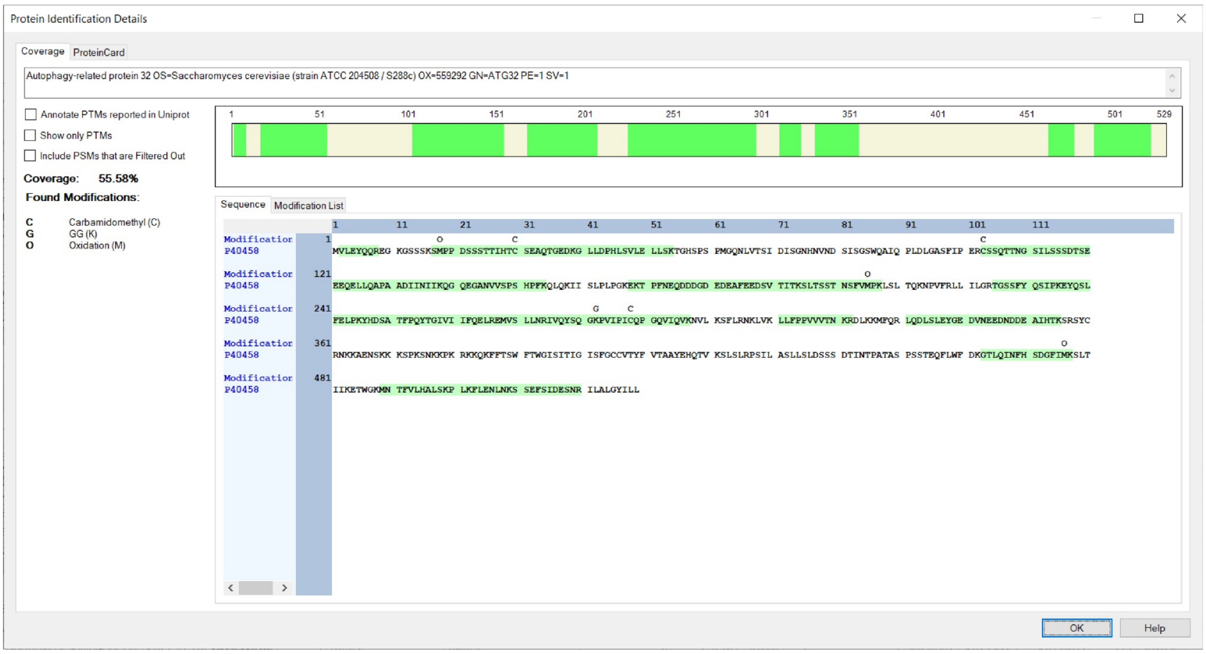

#### Supplementary Table 1

##### STRAINS

| **Souche** | **Genotype** |
|---|---|
| BY4742 + Idp1-GFP+ Atg32-V5 | Mat α ; *his3Δ1;leu2Δ0 ; lys2Δ0 ; ura3Δ0 ; pIDP1-GFP, pYES-Atg32-V5* |
| BY4742 + pPROM-*ATG32-β-galactosidase*, | Mat α ; *his3Δ1;leu2Δ0 ; lys2Δ0 ; ura3Δ0 ;* YEp357-prom*ATG32-lacZ* |
| Δ*atg32* + Idp1-GFP+ Atg32-V5 | BY4742 *atg32*::*kanMX4* p*IDP1-GFP, pYES-Atg32-V5* |
| Δ*atg32* + HA- Atg32-V5 | BY4742 *atg32*::*kanMX4* p*HA-Atg32* |
| Δ*bre5* + Idp1-GFP+ Atg32-V5 | BY4742 *bre5*::*kanMX4* p*IDP1-GFP, pYES-Atg32-V5* |
| Δ*ubp3*+ Idp1-GFP+ Atg32-V5 | BY4742 *ubp3*::*kanMX4* p*IDP1-GFP, pAtg32-V5* |
| Δ*yme1*+ Idp1-GFP+ Atg32-V5 | BY4742 *yme1*::*kanMX4* p*IDP1-GFP, pAtg32-V5* |
| Δ*atg5* + Atg32-V5 | BY4742 *atg5*::*kanMX4 pYES-Atg32-V5* |
| Δ*atg8* + Atg32-V5 | BY4742 *atg8*::*kanMX4, ppYES-Atg32-V5* |
| Δ*atg11*+ Atg32-V5 | BY4742 *atg11*::*kanMX4, pYES-Atg32-V5* |
| Δ*atg32* + Idp1-GFP | BY4742 *atg32*::*kanMX4, pIDP1-GFP* |
| Δ*atg32*+ HA-Atg32 | BY4742 *atg11*::*kanMX4, pHA-Ag32* |
| *pre2-2 + Atg32-V5* | BY4742 *his3D1; leu2D0; met15D0; ura3D0 pre2-2::KanMX4 pYES-Atg32-V5* |
| Δ*atg32*Δ*pho8* + mtPho8 | BY4742 *atg32*::*kanMX4 pho8::HIS3* pFL39-*COXIV-PHO8Δ60* |
